# PicoCam: High-resolution 3D imaging of live animals and preserved specimens

**DOI:** 10.1101/2024.07.16.603742

**Authors:** Josh Medina, Duncan Irschick, Kevin Epperly, David Cuban, Rosalee Elting, Lucas Mansfield, Nora Lee, Ana Melisa Fernandes, Felipe Garzón-Agudelo, Alejandro Rico-Guevara

## Abstract

1. The PicoCam system is a multi-camera photogrammetry rig used for generating high-resolution 3D models of animals in the field, or preserved specimens in laboratory or museum settings. The digital measurement of 3D models is increasingly useful for studying body shape. However, methods that capture sub-millimetric detail often do so at the cost of portability and versatility; this system aims to bridge this gap.

2. The PicoCam system employs 3D digital photogrammetry, a process that generates accurate, full-color, 3D models from sequences of photographs. By using six cameras and a rotating base, the system is able to capture multiple angles in rapid succession – a key advantage for both 3D-imaging live specimens and efficiently scanning museum specimens. Through the use of macro lenses and high-resolution camera sensors, this system can capture sub-millimetric detail without sacrificing portability.

3. In this study, we 3D imaged the bills of 19 species of hummingbirds using the PicoCam system and measured length, height, width, surface area, and volume of their bills. We examined eight species in the field and 11 from the Burke Museum in Seattle. We chose Hummingbird bills as a model system, as their fine-scale interspecific differences in 3D shape can have significant functional and behavioral implications, and could tell us more about how these traits predict habitat and resource use.

4. The prospect of a common 3D-imaging method for both museum and field use is compelling when documenting structures’ shape within and among species. The PicoCam system is also valuable for quantifying the 3D shape of fine-scale phenotypes (like hummingbird bills) that benefit from digital measurement, preservation, and improved accessibility. Lastly, the PicoCam system allows “digital 3D collection”, by which the shape of biological structures in the field can be recorded and stored in a public database without the need to collect a specimen. This opens the door to studies in which multiple 3D image captures of the same individual across different time scales permit 3D shape/color comparisons (e.g., seasonal, ontogenetic changes).

## Intro

The diverse and novel phenotypic adaptations of organisms have inspired generations of researchers to study and categorize biological forms. These forms, and their underlying functions, have occupied the attention of functional morphologists, who compare them among species as a way of identifying adaptations (Lauder, 1981; Losos, 1990; Irschick et al. 1997; Biewener & Patek, 2018). With the development of new tools and methods for quantifying biological form, researchers are able to gather new types of data and ask novel questions about form and function (Rohlf & Marcus, 1993; Zelditch et al., 2012). One of the more recent innovations is the new ability to reconstruct digital 3D models that accurately represent an organism’s body shape while alive (Irschick et al., 2022; Leménager et al., 2022). From the fractal geometry of the nautilus shell to the complex curves of bird ornaments, there is a vast array of complex 3D shapes in nature. These surfaces are difficult to measure using 2D measurements taken with traditional measurement tools like calipers (Cardini, 2014; Buser et al., 2017), or require intensive, landmark-based techniques for deriving 3D coordinates using distance-based methods (Dryden & Mardia, 2016). The need for accurate 3D shape analysis techniques has led to the adoption of 3D-imaging techniques such as computed tomography (CT scanning), laser scanning, and digital photogrammetry. While still comparatively novel, the adoption of digital 3D methods is considered a revolutionary development for the documentation of organismal shape (Adams & Rohlf, 2004; Ziegler et al., 2010; Davies et al., 2017).

The most popular 3D imaging tools fall into distinct niches, each one with its own trade-offs (Díez Díaz et al., 2021; Donato et al., 2020; Mendonca et al., 2013). 3D-imaging tools that preserve sub-millimetric, fine-scale morphology typically trade high resolution for low portability and limited versatility. For example, Micro-CT scanners provide high-resolution images of both external and internal anatomy. However, they aren’t field-portable, have constraints on the shape and size of the objects that can fit inside the machine to be scanned, cannot reconstruct color, and can be prohibitively expensive (Plum & Labonte, 2021; Irschick et al., 2020). Other 3D-imaging tools, such as laser or structured-light scanners, have the opposite problem – portability at the cost of resolution and capture speed. Digital photogrammetry is perhaps the most versatile of these tools, in which objects of varying sizes can be digitized in full color at a range of resolutions based on the camera system in use (Medina et al., 2020). With the use of multiple cameras, it is possible to quickly (i.e., within seconds) capture the data required to reconstruct complex shapes.

Several field-portable photogrammetry systems have been designed to quantify the body shape of live animals in the field. The BeastCam system (Irschick et al., 2020a, b), for instance, is able to make 3D-scans of small (< 50 cm length) live reptiles, amphibians (Irschick et al. 2020a), as well as larger animals such as sea turtles (Irschick et al. 2020b). This method has several advantages, such as the prospect of ‘catch and release’ collection - creating 3D models of live animals that can be measured, stored, and shared in a digital format. These 3D models can be studied without regular access to a museum or field site, using measurements that are impractical to perform on live animals or impossible to make with hand-tools. However, the Beastcam system, as well as other published or commercial field-portable photogrammetry-based tools, do not have the sub-millimetric resolution available in lab-based 3D-imaging tools. High-resolution ‘macro’ photogrammetry systems are possible, such as the ScAnt system (Plum & Labonte, 2021), but are designed for use on mounted arthropods or other small objects, and are not intended for use on live animals in the field. To this end, there is a need for a field-portable system capable of reliably producing full-color high-resolution 3D scans of live organisms that can move or change shape quickly (e.g., wilting in thin-petal flowers).

Herein we present the PicoCam system, a multi-camera photogrammetry tool designed for the 3D-imaging of fine-scale (<0.2 mm resolution) morphology in both museum specimens and live animals in the field. The PicoCam system utilizes macro photography to maximize scan resolution without sacrificing the high-throughput needed to digitize specimens at a range of different sizes and shapes. With a multi-camera setup, morphological features of live animals are scanned via the simultaneous capture of several images from multiple angles. Because the camera mounts are modular, the PicoCam system can be moved and assembled quickly and without tools or external power. Multiple camera mounts allow flexibility in camera positions, lenses, filters, and different camera types according to the needs of the investigator. We demonstrated the utility and flexibility of this system for 3D imaging fine-scale morphological features both in the field for live animals, and in the museum, for preserved specimens.

Using the PicoCam system, we 3D-imaged the bills of 19 species of hummingbirds. This includes eight species of live hummingbirds in the field, and eleven species of preserved hummingbirds from the Burke Museum. Bird bills have been central to conversations of adaptation and biological form/function. For example, variation in bill morphology among bird species represents a textbook example of adaptive radiation (Cooney et al., 2017; Schulter, 2000). Prior work has shown how various bill shapes map onto key dietary functions, such as cracking open seeds, gleaning insects, and extracting nectar. Hummingbirds in particular exhibit tremendously variable bill shapes, in large part because of their mutualistic interactions with plants in which the bill-flower shape/length fit benefits both parties (reviewed by Rico-Guevara et al. 2021). For these birds, fine-scale differences in bill shape can have a large impact on function, such as facilitating access to food resources or affecting performance in intra-sexual combat (Maglianesi et al., 2014; Rico-Guevara & Araya-Salas, 2015, Rico-Guevara et al. 2019). Among or within species variation in height or width, cross-sectional area across the length, surface area, or volume can also potentially alter the function of hummingbird bills. The accurate measurement of fine-scale differences in hummingbird bills represents a fitting case study for 3D imaging tools like the PicoCam system. With high-resolution 3D models generated from both live animals and preserved specimens, we aim to demonstrate a versatile new tool for quantifying organismal shape in both field and museum settings.

## Methods

### 1. Setup

#### 1.1 PicoCam assembly

The PicoCam system consists of six cameras positioned around an aluminum base plate (60cm x 60cm), facing in toward an elevated, rotating platform (30cm x 30cm)(Fig. 1A). Each camera is attached to the platform via an articulating camera arm (Manfrotto, Cassola, Italy) by which its height, angle, and position along the baseplate edge can be adjusted. The platform was designed to support up to twelve arms/cameras. More cameras generally improve the speed of photo capture and the reliability of 3D reconstruction. Subjects were placed on the rotating stage for scanning, and simultaneously photographed by all cameras via remote trigger. The platform rotates to allow for multiple photos to be taken from different angles around the subject, as well as different heights. The apparatus can be set on flat surfaces such as tables with adjustable feet or can be deployed in the field with camera tripods to account for uneven terrain (Fig. 1B). The setup can be arranged for imaging either stationary and revolving subjects; the former is useful for subjects that aren’t suited to the motion of a rotating platform, such as a bird being held in the hand (Fig. 1C).

**Figure 1:**
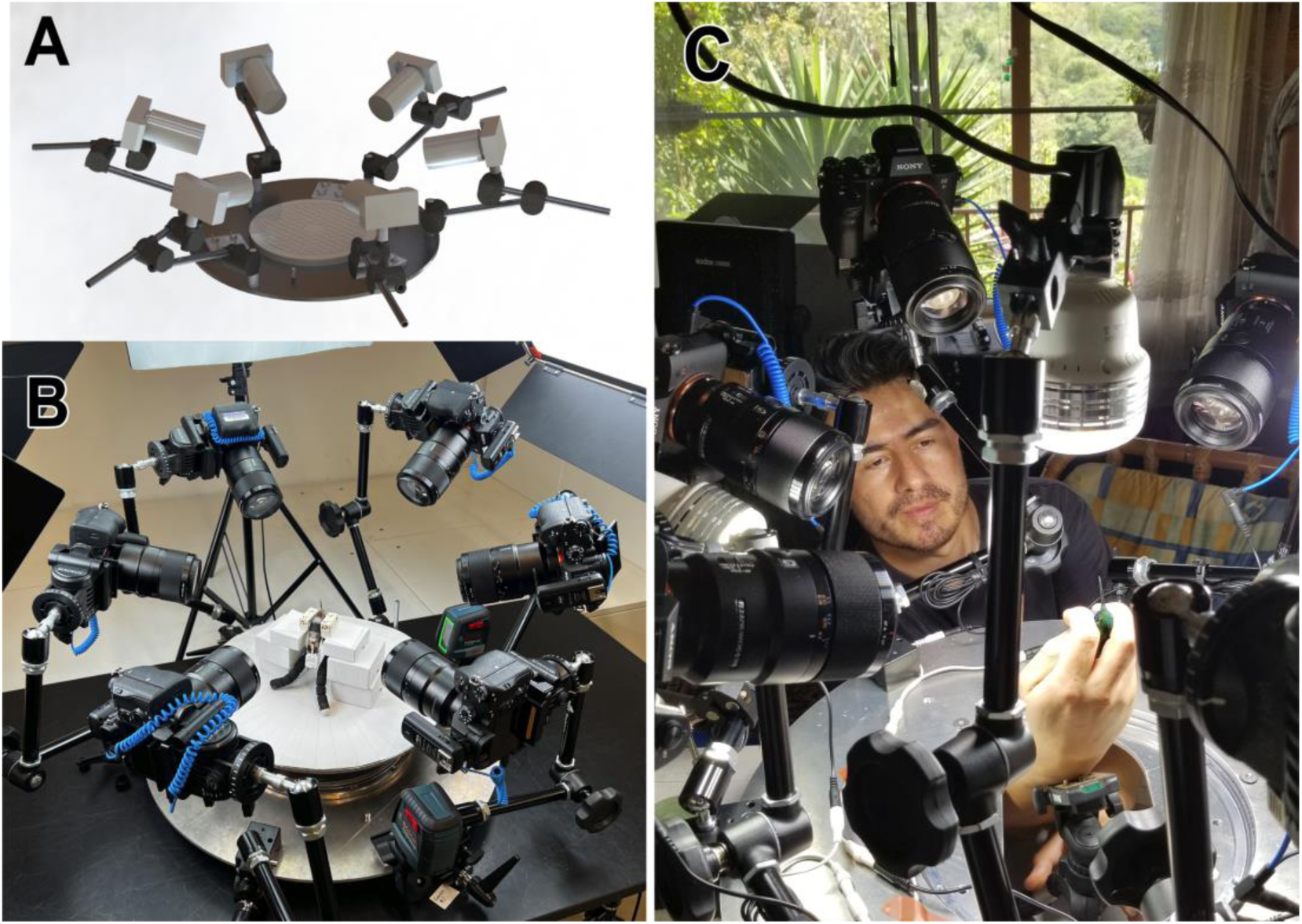
A. A 3D CAD mockup of PicoCam, showing six cameras attached via articulating arms to a metal baseplate, facing a rotating platform that will contain the specimen. B. PicoCam assembled indoors at the Centro Colibri Gorriazul. LED lights have been attached to PicoCam to assist with illumination. C. In this ‘inverted’ setup, the cameras and lights rotate *around* the specimen.

#### 1.2 Cameras, lenses, and lights

Six Sony A7riii mirrorless cameras were attached to the PicoCam system, each equipped with Sony 90mm macro lenses and synchronized via remote trigger (Vello, New York City, United States). Any DSLR or mirrorless camera with a large (24+ megapixel) sensor size could be substituted. The 90mm macro is ideal for capturing the surface detail of small objects like bills of small birds. For larger objects, a 50mm macro lens can be substituted. 90W LED panel lights (SWIT, Santa Clara, United States) were used with adjustable color-temperature set to 5500k. Four panels were oriented around the Picocam system to obtain diffuse, consistent exposure without harsh shadows or glare.

#### 1.3 Case study

The PicoCam system was brought to the Centro de Investigación Colibrí Gorriazul, a research station on the western slope of the Eastern Cordillera in Cundinamarca, Colombia (04° 23’ N, 74° 21’ W, 1,712 m a.s.l.). Eight abundant hummingbird species were chosen: *Anthracothorax nigricollis*, *Amazilia tzacatl*, *Chalybura buffonii*, *Colibri coruscans*, *Colibri cyanotus*, *Colibri delphinae*, *Saucerottia cyanifrons*, and *Phaethornis guy*. One female and one male were captured for each of these species. Most hummingbirds were captured using a Hall trap (a mesh drop net placed around a nectar feeder) (IACUC protocol number PROTO202000143, “4498-05: Nectarivore Feeding, Flight, and Ecology”). *Phaethornis guy* was captured using a 12-m long mist net near Heliconia flowers. Once a hummingbird was captured, we hand-fed it artificial nectar (sugar water, 20%) with an oral syringe to ensure it was well-fed, took standard photographs, and used the Picocam system. Hummingbirds were released shortly after processing. Before being released, the outer surface of the bill was cleaned with tissue to get rid of nectar residue.

#### 1.4 Stand preparation

During live bird processing, we created a hummingbird holding stand to prevent excessive movement, and to minimize environmental stimulus during photography. We used foam blocks that were secured via velcro straps to provide a small area where the bird could be gently held. The foam blocks were attached via magnet to a small GorillaPod tripod, and centered on the rotating platform. For live birds, a cloth with a small hole in the center was gently placed over the bird’s head, obscuring the eyes and exposing the bill from the base. A 3D-printed ‘helmet’ (Fig. 2) was placed over the cloth, onto which were printed coded targets that assist with photo alignment and scale calibration during the photogrammetry process. Once the bird was secured to the stand, the bill was aligned with a vertical straightedge to ensure it was perpendicular to the platform, facing up toward the cameras. To assist with bill alignment and centering, two self-leveling, cross-line lasers (Bosch, Gerlingen, Germany), one red and one green, were attached to the baseplate at perpendicular positions.

**Figure 2:**
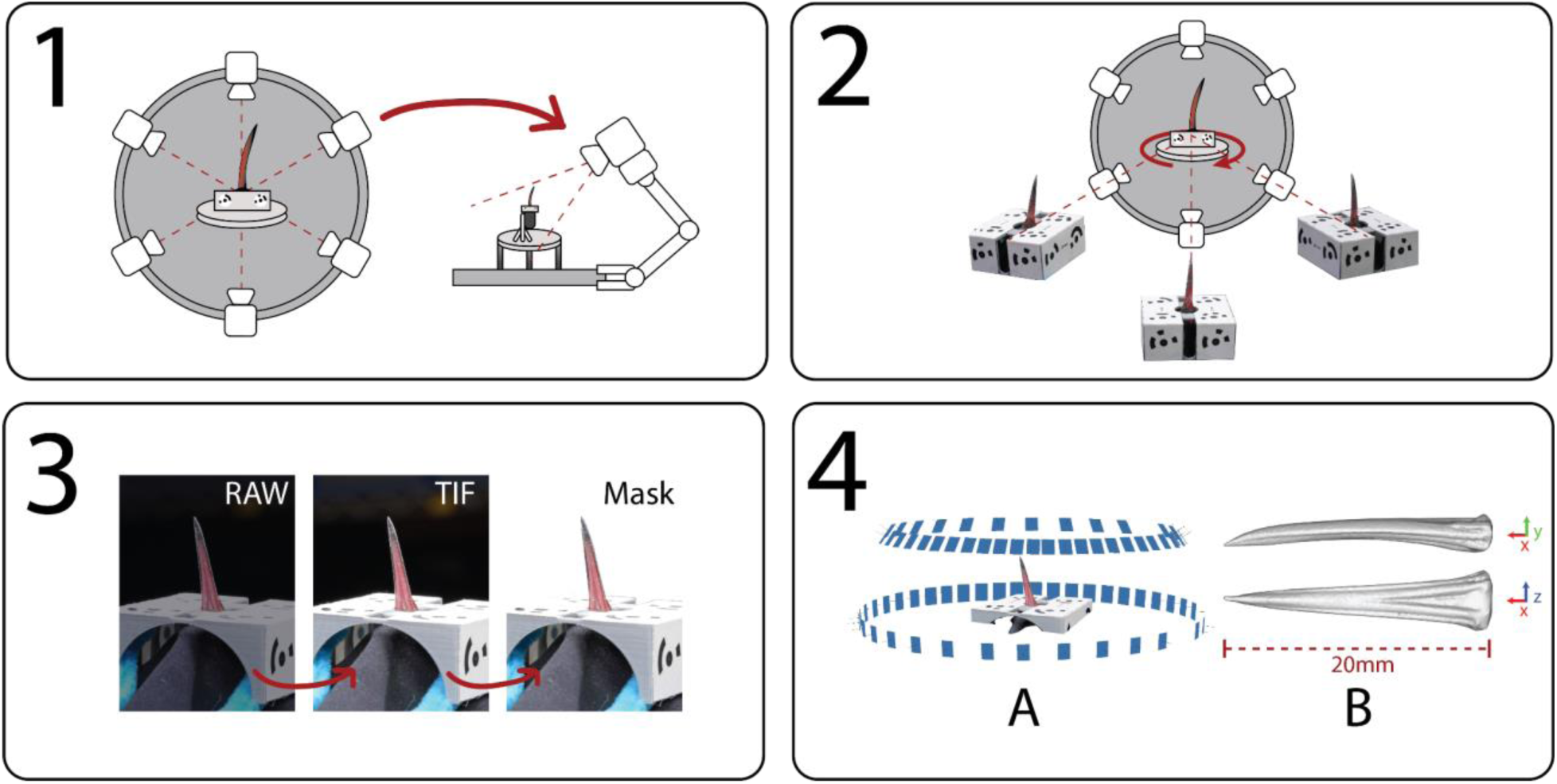
A graphical overview of PicoCam operation. First (1), PicoCam is assembled from six cameras attached via articulating arms to a metal base plate. Above the plate, a rotating platform holds a foam holder containing a hummingbird. 2. During photo capture, each camera simultaneously photographs the hummingbird from different angles. To maximize the number of angles in the scan, the platform is rotated and new photos are taken until either a rotation is completed or the hummingbird shifts position. 3. The RAW photos are adjusted and exported as TIFS. During this process, the backgrounds can be optionally removed via image masks. 4. The photos are aligned using digital photogrammetry software, creating a 3D mesh (A). The 3D mesh can then be separated and analyzed (B) using 3D modeling software.

### 2. Photo capture

#### 2.1 Camera alignment & settings

Camera positions were adjusted to enable maximum overlap between photographs. For capturing hummingbird bills, the cameras were oriented vertically, and staggered at different heights and angles to provide a maximum number of viewpoints. Typically, two cameras would face the bill horizontally, two cameras were raised 4-6 inches and angled down by ∼30° and the last two were raised 6-12 inches and angled down by ∼60°. Camera settings were adjusted to ensure that images were sharp, in-focus, and well exposed. Typical settings used were f22, Shutter Speed of 1/60 and ISO of 2000. During each photo capture session, an X-rite color checker was used to record exposure and color information for image calibration later on. This was particularly important with species with colored bills (e.g., A. tzacatl), but not done for museum specimens because the colors fade after collection.

#### 2.2 Photography

The cameras were triggered via remote trigger to prevent camera shake. After the cameras were triggered, the platform was rotated (by ∼10° for live birds and ∼4.5° for museum specimens) and a set of new photos were taken. This process was repeated until a complete rotation was made. The ideal number of photos depends on the object shape and the desired resolution of the scan (Fig. 3). With live birds, it was useful to take as many photos as possible before the bird moved. Minor shifts in position were enough to reset the process. After several iterations of our bird stand, we were able to keep small movements to a minimum and allow for 100 - 480 photos per scan. After the desired number of photos were taken (and/or rotations have been made) the specimen was removed from the stage. The bill was measured with digital calipers, photographed from standard angles, and then the bird was released. Processing times for live and preserved specimens ranged from about 10-20 minutes.

**Figure 3.**
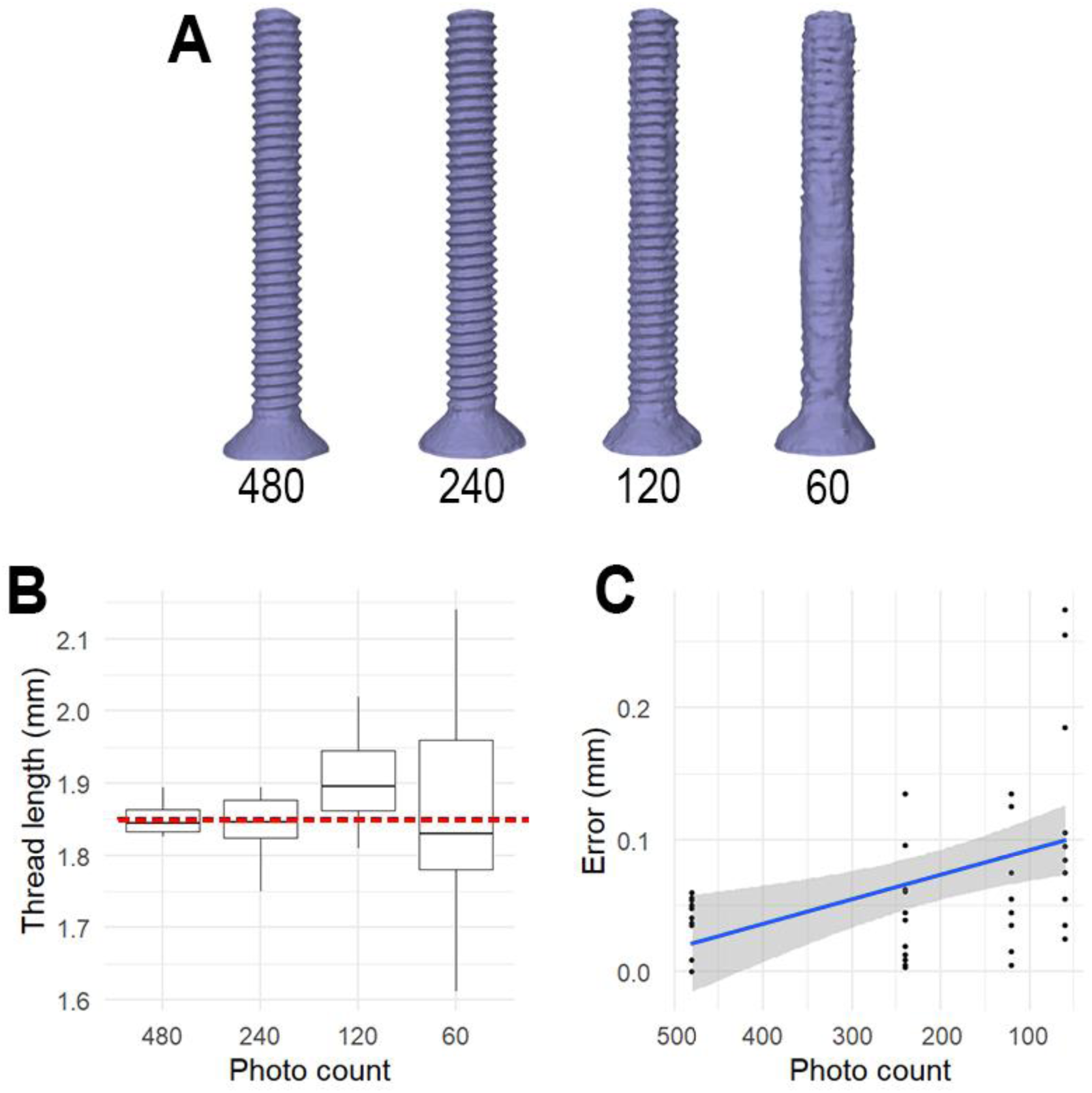
To test the effects of photo count on 3D resolution, we 3D-imaged the same reference screw (M4 - 0.7×16mm) four times using different photo counts. The resulting 3D meshes (A) show decreasing resolution that begins to affect the number of visible threads somewhere between 60 and 120 photos. The width of the threads of each screw model was measured and compared (B), with the red dotted line representing the average ‘real’ thread width measured on the physical screw. The difference between the ‘real’ thread width and the average of each 3D mesh was measured as error (C) and compared. The relationship between measurement error and photo count can be read from the blue trend line.

### 3. Image processing

#### 3.1 Color + exposure correction

Images were saved in RAW format and loaded onto a computer using a Multi-SD card reader. Camera settings were adjusted to add prefixes to each file specifying which camera (1-6) they were taken from, along with accurate time and date. RAW files were imported into Adobe Lightroom for processing. Photos with obvious visual defects (blurry, out of focus, etc) were removed. Color, white-balance and exposure were corrected according to color reference photos taken just prior to photo capture. Images were exported in 16-bit TIFF format, which provided better detail than lossy alternatives (like JPGs) during 3D reconstruction. With the museum specimens, lighting conditions and white balance were calibrated in each camera ahead of time, allowing this step to be skipped entirely.

#### 3.2 Background masking

For some subjects, we used image masks to remove the background of images to remove extraneous details that interfered with image alignment (see *mesh processing*). In these cases, Adobe Photoshop (Adobe, https://www.adobe.com/) was used to automatically batch-process large numbers of image masks. For the bird bills, the inclusion of coded targets improved image alignment to the extent that image masks were no longer necessary.

### 4. Mesh generation

#### 4.1 Photo alignment

Processed TIFF images were imported into Agisoft Metashape version 2.1.1 (Agisoft, www.agisoft.com) a photogrammetry software package that creates 3D models out of 2D images. After the images were loaded, the software located coded targets present in the photos which assist in photo alignment. 12 coded targets were located at fixed positions around the specimen stand, oriented along the X, Y, and Z axes to minimize distortion (see Figure 2.) During photo alignment, a 3D point cloud was generated and extraneous details (such as the stand or background) were removed. Images that failed to align were either removed or corrected. The coded targets used in photo alignment were then re-used for scale calibration, with the centers of pairs of targets set a known distance (20mm) apart on the specimen stand.

#### 4.2 Mesh processing

After scale calibration, a 3D polygonal mesh was generated from the point cloud, with a resolution in the range of between 100-300,000 polygons. This mesh was exported in .OBJ format, and loaded into Blender version 4.1.0, an open-source 3D modeling program (Blender Foundation, www.blender.org). The bill was then isolated and any topological defects from mesh generation were corrected. The mesh was made ‘watertight’ by filling the hollow base of the bill, along with any other holes in the surface. This ensures the mesh is one continuous closed surface, which is important for volumetric measurements. The edited meshes were exported as an .OBJ file for use in other software, or used in Blender for basic shape analysis.

### 5. Analysis

#### 5.1 Bill measurements

In total, bills representing 19 species of hummingbird were 3D-imaged and measured. This includes eight species of live birds measured in the field and 11 species from museum specimens (Fig. 4). All bills were first measured for bill length, width, and height using calipers. These measurements were then repeated using 3D models in Blender. Length measurements were limited to the exposed culmen, measured from the point where feathers begin at the upper ridge to the tip of the bill. Width and height were taken at the same position at the base of the bill. The bills were then measured for surface area, and volume using Blender’s built-in 3D printing toolbox. For volumetric measurements, the portion of the bill from the point where feathers begin to the tip was isolated. All bills measured this way were closed, with the upper and lower mandibles forming one continuous 3D surface.

**Figure 4.**
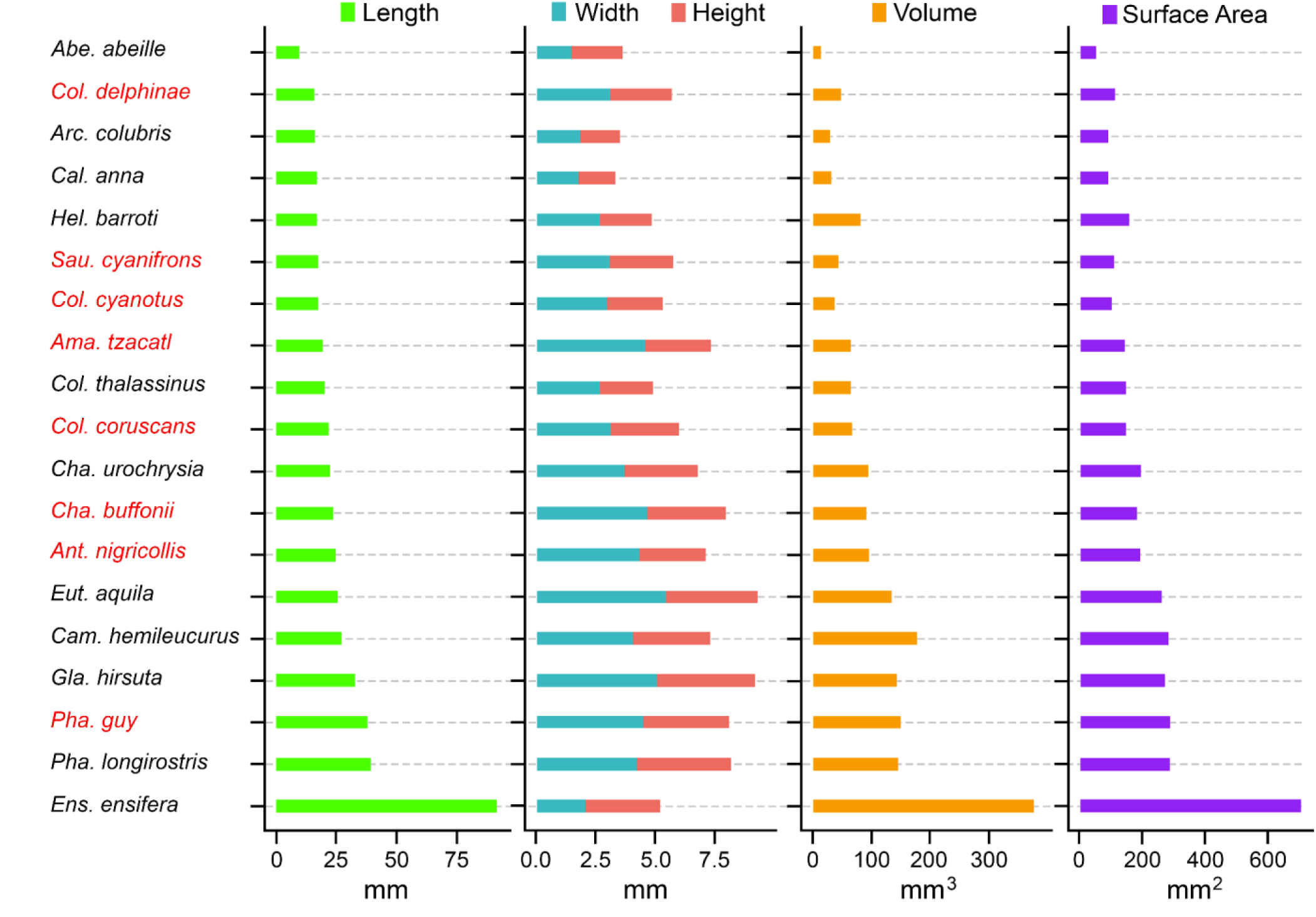
A visual overview comparing bill dimensions derived from 3D models representing 19 species of hummingbirds. Metrics include Length, Width and Height (mm) along with Volume (mm^3^) and Surface Area (mm^2^). The names of 3D models derived from live birds are marked in red. A comprehensive list of species is available in the supplementary material.

#### 5.2 Phylogeny

A time-calibrated phylogenetic tree for hummingbirds, originally from McGuire et al. (2014) and modified to include species as in Colwell et al. (2023), was pruned to include the 19 species in the dataset (Fig. 5). The resulting phylogeny was then used to test for phylogenetic signal using phylogenetic cladograms and Moran’s I index. Next, several tests examined which combinations of basic dimensions (length, width, and height) best predicted surface area, and volume. Phylogenetic Generalized Least Squares (PGLS) was used to examine the relationship among volume and the three different dimension traits while accounting for phylogenetic non-independence. The model was fit using the Maximum Likelihood estimate of λ (Brownian motion). Due to the number of predictor variables, the results were compared using ANOVA tables. All analyses were conducted in r version 4.4.1 (R Core Team, 2013), using phytools (Revell, 2012) to generate the phylogenetic tree and caper (Oem et al., 2013) to perform the PGLS.

**Figure 5.**
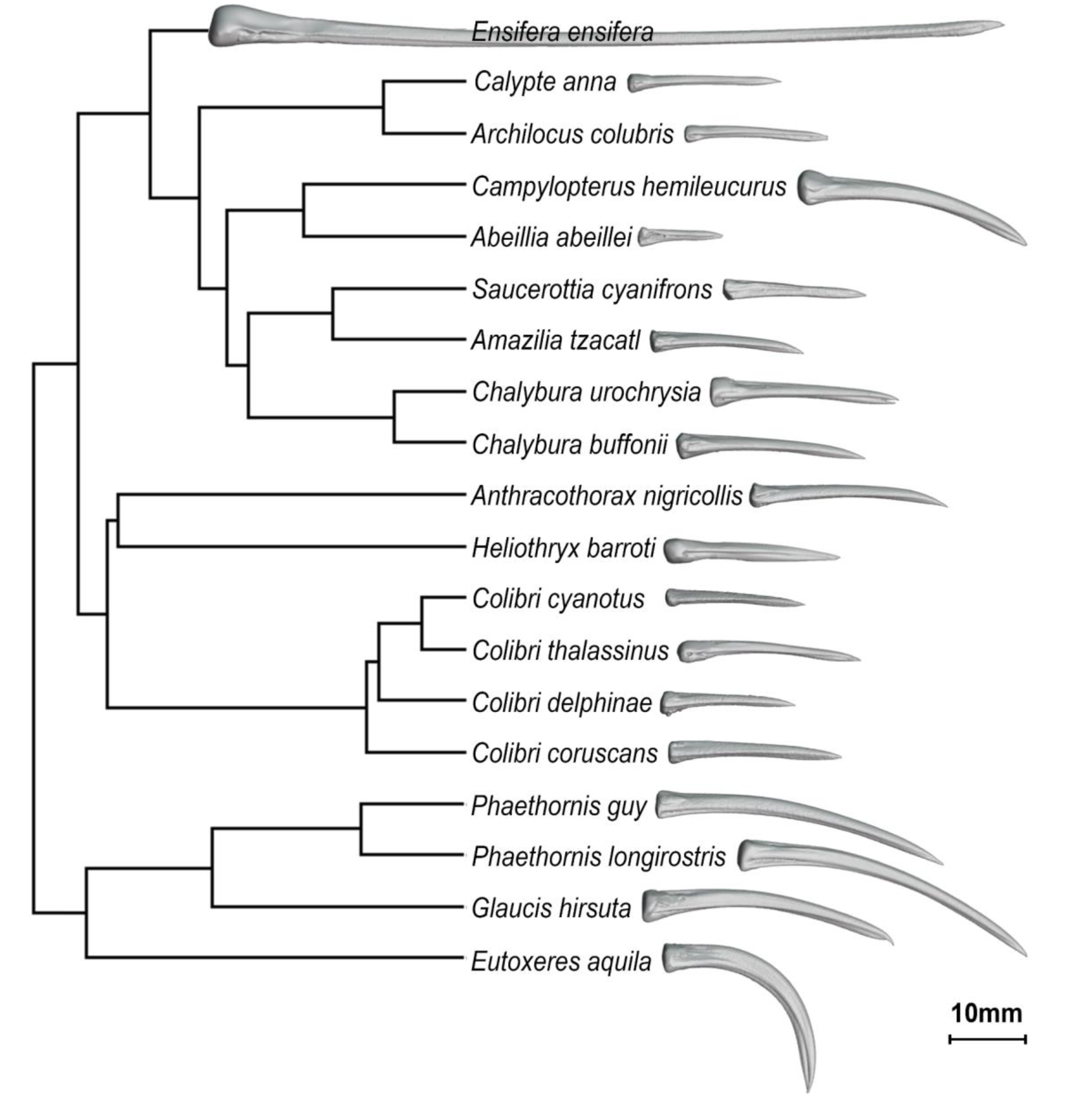
A phylogenetic tree showing 19 species of hummingbird. Each species has a corresponding 3D bill model generated by PicoCam, shown in profile view. The tree include species from 5 different clades, from top to bottom: Brilliants (Ensifera), Bees (Calypte + Archilocus), Emeralds (Campylopterus to Chalybura), Mangoes (Anthracothorax to Colibri), and Hermits

## Results

Most bill measurements varied extensively between species (Fig. 4), except for height and width. The longest bill (*Ensifera ensifera*) was 9.4x longer than the shortest bill (*Abeillia abeillei*). The same birds had less than a 0.4x (40%) difference in bill height, illustrating a general trend of bill length being more variable among species than width or height (Fig. 4). A full list of specimens, along with measurement data, is available in the supplementary material. The 3D models generated for this study are available for viewing and download via sketchfab (www.sketchfab.com) [https://sketchfab.com/irschicklab].

The accuracy of the models was verified by comparing the digital measurements to physical equivalents. Measurements of exposed culmen length, bill height, and bill width were taken using calipers on both live bird bills and museum specimens. The average error was 0.432mm (SD 0.32) for exposed culmen length, 0.355mm for bill width (SD 0.30), and 0.36mm for bill height (SSD 0.299). Further assessments of accuracy, including the effects of photo count on 3D resolution, were made by measuring the threads of a 16mm reference screw (Fig. 3).

When testing for autocorrelation, no traits scored a Moran’s I Index above or below 2, with the exception of *Ensifera ensifera*. Phylogenetic correlograms also returned no significant signs of phylogenetic signal affecting our dataset. The PGLS results show that bill length was significantly correlated with volume (PGLS: F = 447, p < 0.001, λ = 0.47), as well as bill width (PGLS: F = 64, p < 0.001, λ = 0.47). Height’s correlation with volume was also significant (p < 0.01). Interaction between length, width, and/or height did not significantly predict volume. Looking at the model coefficients, there was also a positive relationship between length and volume, but it was not significant (PGLS: slope ± SE = 2.261 ± 0.763, t = 2.96, df = 10, p = 0.014, λ = 0.47). The PGLS results showed similar trends for surface area. Length significantly correlated with surface area (PGLS: F = 754, p < 0.001, λ = 0.71), as did Width. (PGLS: F = 43, p < 0.001, λ = 0.71). Height also correlated with surface area (p < 0.01). As before, interaction between dimension variables did not significantly correlate with surface area. There was also a positive, but not significant, relationship between length and surface area. (PGLS: slope ± SE = 1.51 ± 0.45, t = 3.4, df = 10, p = 0.006, λ = 0.71).

## Discussion

In this study, we described a new device (the PicoCam system), which allowed us to 3D-image the bills of 19 species of hummingbirds, eight of which were generated from live birds in the field, and 11 from museum specimens. From the resulting 3D models, we were able to derive height, width, length, surface area, and volume data. The ability to create high-resolution 3D models of both live animals in the field and museum specimens is valuable for several reasons, which we explain below.

### Challenges and value of photogrammetry for bird bills

One of the most significant challenges for anatomists and functional morphologists is to accurately visualize detailed anatomical structures. Whereas this challenge can be fairly straightforwardly met using CT scanning on preserved specimens, many subtle morphological features are altered during various kinds of preservation processes, and there is a value in quantifying these forms in live animals in the field. Hummingbird bills represent a good example of this problem. Their bills, like those of other birds (Cooney et al., 2017), are challenging for 3D photogrammetric reconstruction because their elongated and delicate structures are shiny, and lack obvious landmarks.

Bird bill shape variation, both interspecific and intraspecific, has provided demonstrations of coevolution (Stiles, 1981), ornamentation (Romero-Diaz, 2022), and weaponization (Rico-Guevara & Araya-Salas, 2015) – sometimes all at once (Rico-Guevara et al., 2019). With measurements such as surface area, curvature, or sharpness, inferences can be made about ecological niche, such as foraging strategy (Rico-Guevara et al., 2019, 2021). This may help researchers evaluate ecological tradeoffs mediated by functional variation, such as between bite force and the ability to manipulate complex objects. Importantly, these trade-offs could be examined both among individuals, as the Picocam system can be potentially used to scan large numbers of individuals over the course of the field season, or among species.

### The ‘digital specimen’

Given that some characteristics of organisms can change after collection, such as the 3D shape of flowers in herbaria, or the bill color in some birds, the Picocam system could generate life-like (sensu Irschick et al. 2022) models of museum specimens that can be linked digitally with the preserved specimens. Although we recognize the irreplaceable value of collecting organisms for museum collections (e.g., Nachman et al. 2023), the PicoCam system also allows for a ‘catch-and-release’ approach to 3D data gathering, in which minimal scan times and rapid release allows researchers to non-invasively gather organismal shape data. This would make it easier to collect specimen data in regions where collecting, transporting, or obtaining permits for specimens is difficult or impossible. These issues are of increasing relevance to researchers aiming to gather data from live specimens globally. Further, the PicoCam system could allow researchers to scan and quantify structures from individuals who would regularly not be candidates for scanning due to life history (e.g., females who could be nesting) or conservation status (e.g., endangered taxa that could not regularly be collected). This method of collection and sharing across cloud-based servers allows for the extension of the intellectual properties from these scans to communities beyond those with resources to visit museums in the global north.

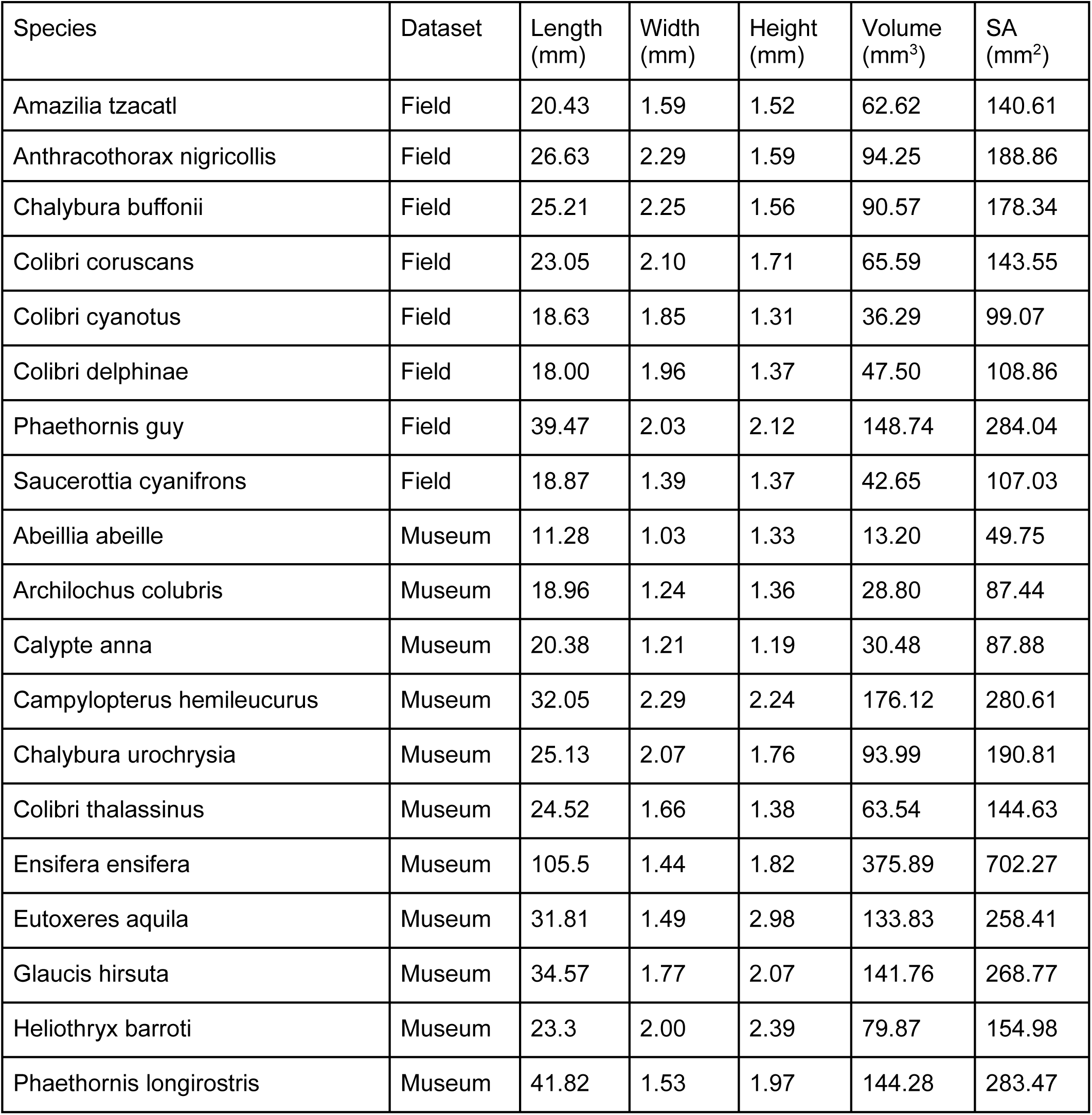

## References

Adams, D. C., Rohlf, F. J., & Slice, D. E. (2004). Geometric morphometrics: ten years of progress following the ‘revolution’. Italian journal of zoology, 71(1), 5–16. 10.1080/11250000409356545

Biewener, A., & Patek, S. (2018). Animal locomotion. Oxford University Press.

Bot, J. A., Irschick, D. J., Grayburn, J., Lischer-Katz, Z., Golubiewski-Davis, K., & Ikeshoji-Orlati, V. (2019). Using 3D photogrammetry to create open-access models of live animals: 2D and 3D software solutions. Grayburn et al., eds. D, 3, 54–72.

Buser, T. J., Sidlauskas, B. L., & Summers, A. P. (2018). 2D or not 2D? Testing the utility of 2D vs. 3D landmark data in geometric morphometrics of the sculpin subfamily Oligocottinae (Pisces; Cottoidea). The Anatomical Record, 301(5), 806–818. 10.1002/ar.23752

Cardini, A. (2014). Missing the third dimension in geometric morphometrics: how to assess if 2D images really are a good proxy for 3D structures?. Hystrix, 25, 73–81. 10.4404/hystrix-25.2-10993

Colwell, R. K., Rangel, T. F., Fučiková, K., Sustaita, D., Yanega, G. M., & Rico-Guevara, A. (2023). Repeated evolution of unorthodox feeding styles drives a negative correlation between foot size and bill length in hummingbirds. 10.1086/726036

Cooney, C. R., Bright, J. A., Capp, E. J., Chira, A. M., Hughes, E. C., Moody, C. J., … & Thomas, G. H. (2017). Mega-evolutionary dynamics of the adaptive radiation of birds. Nature, 542(7641), 344–347. 10.1038/nature21074

Davies, T. G., Rahman, I. A., Lautenschlager, S., Cunningham, J. A., Asher, R. J., Barrett, P. M., … & Donoghue, P. C. (2017). Open data and digital morphology. Proceedings of the Royal Society B: Biological Sciences, 284(1852), 20170194. 10.1098/rspb.2017.0194

Díez Díaz, V., Mallison, H., Asbach, P., Schwarz, D., & Blanco, A. (2021). Comparing surface digitization techniques in palaeontology using visual perceptual metrics and distance computations between 3D meshes. Palaeontology, 64(2), 179–202. 10.1111/pala.12518

Donato, L., Cecchi, R., Goldoni, M., & Ubelaker, D. H. (2020). Photogrammetry vs CT Scan: Evaluation of Accuracy of a Low-Cost Three-Dimensional Acquisition Method for Forensic Facial Approximation. Journal of Forensic Sciences, 65(4), 1260–1265. 10.1111/1556-4029.14319

Dryden, I. L., & Mardia, K. V. (2016). Statistical shape analysis: with applications in R (Vol. 995). John Wiley & Sons. iDigBio (2018) https://www.idigbio.org/portal

Irschick, D. J., Vitt, L. J., Zani, P. A., & Losos, J. B. (1997). A comparison of evolutionary radiations in mainland and Caribbean Anolis lizards. Ecology, 78(7), 2191–2203.

Irschick DJ, Bot J, Brooks A, Bresette M, Fossette S, Gleiss A, Gutierrez R, Manire C, Merigo C, Martin J, Pereira M, Whiting S, Wyneken J. (2020a). Using 3D photogrammetry to create accurate 3D models of sea turtle species as digital voucher specimens. Herpetological Review. 51:709–715.

Irschick DJ, Corriveau Z, Mayhan T, Siler C, Mandica M, Gamble T, Martin J, Bot J, Zotos S. (2020b). Devices and Methods for Rapid 3D Photo-Capture and Photogrammetry of Small Reptiles and Amphibians in the Laboratory and the Field. Herpetological Review. 51:716–725.

Irschick, D. J., Christiansen, F., Hammerschlag, N., Martin, J., Madsen, P. T., Wyneken, J., … & Lauder, G. (2022). 3D visualization processes for recreating and studying organismal form. IScience, 25(9).

Lauder, G. V. (1981). Form and function: structural analysis in evolutionary morphology. Paleobiology, 7(4), 430–442. 10.1017/S0094837300025495

Leménager, M., Burkiewicz, J., Schoen, D. J., & Joly, S. (2023). Studying flowers in 3D using photogrammetry. New Phytologist, 237(5), 1922–1933. 10.1111/nph.18553

Losos, J. B. (1990). Ecomorphology, performance capability, and scaling of West Indian Anolis lizards: an evolutionary analysis. Ecological Monographs, 60(3), 369–388. 10.2307/1943062

Maglianesi, M. A., Blüthgen, N., Böhning-Gaese, K., & Schleuning, M. (2014). Morphological traits determine specialization and resource use in plant–hummingbird networks in the neotropics. Ecology, 95(12), 3325–3334. 10.1890/13-2261.1

McGuire, J. A., Witt, C. C., Remsen, J. V., Corl, A., Rabosky, D. L., Altshuler, D. L., & Dudley, R. (2014). Molecular phylogenetics and the diversification of hummingbirds. Current Biology, 24(8), 910–916. 10.1016/j.cub.2014.03.016

Medina, J. J., Maley, J. M., Sannapareddy, S., Medina, N. N., Gilman, C. M., & McCormack, J. E. (2020). A rapid and cost-effective pipeline for digitization of museum specimens with 3D photogrammetry. PLoS One, 15(8), e0236417. 10.1371/journal.pone.0236417

Mendonca, D. A., Naidoo, S. D., Skolnick, G., Skladman, R., & Woo, A. S. (2013). Comparative study of cranial anthropometric measurement by traditional calipers to computed tomography and three-dimensional photogrammetry. Journal of Craniofacial Surgery, 24(4), 1106–1110. DOI: 10.1097/SCS.0b013e31828dcdcb

Nachman, M. W., Beckman, E. J., Bowie, R. C., Cicero, C., Conroy, C. J., Dudley, R., … & Zink, R. M. (2023). Specimen collection is essential for modern science. Plos Biology, 21(11), e3002318. 10.1371/journal.pbio.3002318

Nelson, G., & Ellis, S. (2019). The history and impact of digitization and digital data mobilization on biodiversity research. Philosophical Transactions of the Royal Society B, 374(1763), 20170391. 10.1098/rstb.2017.0391

Orme, D., Freckleton, R., Thomas, G., Petzoldt, T., Fritz, S., Isaac, N., & Pearse, W. (2013). The caper package: comparative analysis of phylogenetics and evolution in R. R package version, 5(2), 1–36.

Page, L. M., MacFadden, B. J., Fortes, J. A., Soltis, P. S., & Riccardi, G. (2015). Digitization of biodiversity collections reveals biggest data on biodiversity. BioScience, 65(9), 841–842. 10.1093/biosci/biv104

Plum, F., & Labonte, D. (2021). scAnt—an open-source platform for the creation of 3D models of arthropods (and other small objects). PeerJ, 9, e11155. 10.7717/peerj.11155

R Core Team, R. (2013). R: A language and environment for statistical computing. http://www.R-project.org

Revell, L. J. (2012). phytools: an R package for phylogenetic comparative biology (and other things). Methods in ecology and evolution, (2), 217-223. doi:10.1111/j.2041-210X.2011.00169.x.

Rico-Guevara, A., & Araya-Salas, M. (2015). Bills as daggers? A test for sexually dimorphic weapons in a lekking hummingbird. Behavioral Ecology, 26(1), 21–29. 10.1093/beheco/aru182

Rico-Guevara, A., Rubega, M. A., Hurme, K. J., & Dudley, R. (2019). Shifting paradigms in the mechanics of nectar extraction and hummingbird bill morphology. Integrative Organismal Biology, 1(1), oby006. DOI:10.1093/iob/oby006

Rico-Guevara, A., Hurme, K. J., Elting, R., & Russell, A. L. (2021). Bene “fit” assessment in pollination coevolution: mechanistic perspectives on hummingbird bill–flower matching. Integrative and Comparative Biology, 61(2), 681–695. 10.1093/icb/icab111

Rohlf, F. J., & Marcus, L. F. (1993). A revolution morphometrics. Trends in ecology & evolution, 8(4), 129–132. 10.1016/0169-5347(93)90024-J

Romero-Diaz, C., Silva, P. A., Soares, M. C., Cardoso, G. C., & Trigo, S. (2022). Oestradiol reduces female bill colour in a mutually ornamented bird. Proceedings of the Royal Society B, 289(1984), 20221677. 10.1098/rspb.2022.1677

Page, Lawrence M., et al. “Digitization of biodiversity collections reveals biggest data on biodiversity.” BioScience 65.9 (2015): 841–842. 10.1093/biosci/biv104

Schluter, D. (2000). The ecology of adaptive radiation. OUP Oxford.

Stiles, F. G. (1981). Geographical aspects of bird-flower coevolution, with particular reference to Central America. Annals of the Missouri Botanical Garden, 323–351. 10.2307/2398801

Ströbel, B., Schmelzle, S., Blüthgen, N., & Heethoff, M. (2018). An automated device for the digitization and 3D modelling of insects, combining extended-depth-of-field and all-side multi-view imaging. ZooKeys, (759), 1. doi: 10.3897/zookeys.759.24584

Zelditch, M., Swiderski, D., & Sheets, H. D. (2012). Geometric morphometrics for biologists: a primer. academic press.

Ziegler, A., Ogurreck, M., Steinke, T., Beckmann, F., Prohaska, S., & Ziegler, A. (2010). Opportunities and challenges for digital morphology. Biology direct, 5(1), 1–9. 10.1186/1745-6150-5-45

